# Genomic Surveillance for Antimicrobial Resistance in *Mannheimia haemolytica* Using Nanopore Single Molecule Sequencing Technology

**DOI:** 10.1101/395087

**Authors:** Alexander Lim, Bryan Naidenov, Haley Bates, Karyn Willyerd, Timothy Snider, Matthew Brian Couger, Charles Chen, Akhilesh Ramachandran

**Affiliations:** Department of Biochemistry and Molecular Biology, Oklahoma State University, 246 Noble Research Center, Stillwater, OK 74078; Center for Veterinary Health Sciences, Oklahoma Animal Disease Diagnostic Laboratory, 1950 W. Farm Road, Stillwater, OK 74078; Department of Microbiology and Molecular Genetics, Oklahoma State University, 307 Life Sciences East, Stillwater, OK 74078

## Abstract

Disruptive innovations in long-range, cost-effective direct template nucleic acid sequencing are transforming clinical and diagnostic medicine. A multidrug resistant strain and a pan-susceptible strain of *Mannheimia haemolytica*, isolated from pneumonic bovine lung samples, were respectively sequenced at 146x and 111x coverage with Oxford Nanopore Technologies MinION. *De novo* assembly produced a complete genome for the non-resistant strain and a nearly complete assembly for the drug resistant strain. Functional annotation using RAST (Rapid Annotations using Subsystems Technology), CARD (Comprehensive Antibiotic Resistance Database) and ResFinder databases identified genes conferring resistance to different classes of antibiotics including beta lactams, tetracyclines, lincosamides, phenicols, aminoglycosides, sulfonamides and macrolides. Antibiotic resistance phenotypes of the *M. haemolytica* strains were confirmed with minimum inhibitory concentration (MIC) assays. The sequencing capacity of highly portable MinION devices was verified by sub-sampling sequencing reads; potential for antimicrobial resistance determined by identification of resistance genes in the draft assemblies with as little as 5,437 MinION reads corresponded to all classes of MIC assays. The resulting quality assemblies and AMR gene annotation highlight efficiency of ultra long-read, whole-genome sequencing (WGS) as a valuable tool in diagnostic veterinary medicine.

## Introduction

Emergence of antimicrobial resistance (AMR) among the most important bacterial pathogens is recognized as a major public health concern. Not only has AMR emerged in hospital environments, it is often identified in community settings, in livestock feedlots and in aquaculture and crop production, suggesting an ever-increasing range of reservoirs of antibiotic-resistant bacteria (1-4). Bacterial response to the antibiotic “attack” is the prime example of genetic adaptation through the interplay of immense genetic plasticity, ranging from mutational adaptations and acquisition of genetic material to alteration of gene expression with fitness consequence of the pathogen (5). As a result, understanding the genetic basis of resistance is of paramount importance to design strategies to curtail the emergence and spread of AMR, as well as to devise innovative therapeutic approaches against multidrug-resistant organisms (6).

With an increased population of pet and farm animals there is a significant risk of humans acquiring drug resistant bacteria from animal sources. One of the major deficiencies in veterinary medicine is the lack of validated data to determine minimum inhibitory concentration (MIC) breakpoints for drug-microbe-host combinations, based on which scientifically sound interpretations can be made regarding whether a pathogen is susceptible or resistant to a specific drug. This is especially true for anaerobic bacteria. The large diversity of domestic and exotic animal species and their numerous microbial pathogens pose significant challenges to developing reliable interpretation criteria for all antimicrobial drugs.

Recently, whole-genome sequencing (WGS) has become an invaluable tool to combat antibiotic resistance (7-12). In addition to the bench-top sequencing platforms like Illumina machines, in 2014 Oxford Nanopore Technologies (ONT) released a portable MinION device through an early access program (the MinION Access Program). MinION devices can be powered by a standard USB3 port and the base-calling step that turns electrical signals into nucleotides is enabled by a cloud-computing platform (13), providing the portability needed for various clinical microbiology settings, especially for political or logistical reasons (14-16). The landmark case for diagnostic investigation of pathogen genomes was the Ebola outbreak in Guinea (17). Using three MinION devices and four laptop computers, 148 sequencing runs were conducted to cover 142 samples, and a rapid turnaround was demonstrated that sequencing process could be performed within an hour with the whole work-flow from amplification, library preparation and sequencing runs brought to completion in 24 hours (18). Currently, sequencing capacity of MinION has reached about 450 bases per second; as the throughput and speed increases, ONT platform might become suitable for real-time pathogen surveillance and clinical diagnostic applications (19).

In veterinary medicine, approaches such as mass spectrometry and multiplexed PCR are commonly used for diagnostic applications. WGS, however, has yet to be adopted as a part of routine diagnostic procedures. In this report we describe the use of WGS for evaluating AMR of *Mannheimia haemolytica,* an important bacterial pathogen of cattle causing bovine respiratory disease (BRD). The annual loss to the cattle industry from BRD is estimated to be around one billion USD in North America (20). Increase in antimicrobial resistant *M. haemolytica* over the years has been recognized (21, 22); and, high prevalence of multidrug resistance of *M. haemolytica* that underscores critical challenges regarding treatments and management practices was also strongly reckoned in a recent study (23). In this manuscript WGS-based detection of AMR of two *Mannheima* strains was determined using ONT’s MinION devices; with the assemblies constructed with as little as 5,347 ONT ultra-long reads, potential bacterial AMR was supported by all genome annotation databases and the genetic AMR of these strains was further confirmed with MIC assays. Using this comparative study as an example, the feasibility and effectiveness of integrating ONT’s MinION technology in clinical investigations was explored.

## Materials and Methods

### DNA extraction for bacterial isolates

*M. haemolytica* OADDL-1 was isolated from the lung of a 3-year-old bull and *M. haemolytica* OADDL-2 was isolated from the lung of a calf of unknown age. Both animals from which the bacterial isolates were obtained had a history of sudden death and a histologic diagnosis of severe acute fibrinous pleuropneumonia. The calf from which the resistant *M. haemolytica* isolate was obtained also had a history of antimicrobial treatment with Tilmicosin and Enrofloxacin. Bacterial isolates were cultured on 5% sheep blood agar (Hardy Diagnostics); five to 10 colonies of bacteria were carefully selected to avoid agar contamination and then suspended in Tris EDTA buffer. Genomic DNA was extracted using the OMEGA Bio-tek EZNA® bacterial DNA kit D3350-01 protocol (OMEGA Bio-tek, Norcross, GA).

### MIC (minimum inhibitory concentration)

Antimicrobial susceptibility of *M. haemolytica* isolates was determined by MIC method on the Sensititre automated system (Thermo Scientific, Waltham, MA, USA), using the Bovine/Porcine panel (BOPO 6F, Thermo Scientific, Waltham, MA, USA) following manufacturer recommendations. Interpretations regarding susceptibility or resistance to antimicrobial were made based on CLSI VET01S guidelines (24).

### MinION Library Preparation and Barcoding

Library preparation was performed following the procedures outlined for SQK-LSK208 sequencing kit and barcoded for multiplexing using the complimentary EXP-NDB002 barcoding kit (Oxford Nanopore Technologies, United Kingdom) with the following protocol adjustments. A total of 1.5 µg of gDNA from each bacterial isolate was sheared in g-tubes (Covaris) at 4200 RPM for a targeted fragment size of 20 Kb. End-repair was performed following the recommended protocol of the manufacturer for Ultra II End-prep enzyme mix (NEB). Adapter ligation reaction incubations were increased to 15 minutes. Bead clean-ups used 0.4x AMPureXP beads (Beckman Coulter, Brea, CA) for additional size selection and elutions were performed at 37^°^C for 20 minutes. DNA concentration of the library was quantified using Quant-IT PicoGreen^®^ dsDNA Assay Kit (ThermoFisher Scientific), measured on Synergy H1, hybrid multi-mode microplate reader (BioTek), confirming final library yields above the recommended 200 ng. In total, two MinION barcoded libraries were prepared separately for sequencing on two flow cells (listed as Library A and B in Table 2).

**Table 1.** Phenotypic susceptibility testing results for bacterial strains *M. haemolytica* OADDL-1 and *M. haemolytica* OADDL-2.

| CLSI Group | Antibacterial Name | Antibacterial Class | *M. haemolytica*<br>OADDL-2 | *M. haemolytica*<br>OADDL-1 |
| --- | --- | --- | --- | --- |
| A | Spectinomycin | Aminocyclitol | **R (> 64)** | **S (= 16)** |
| A | Ceftiofur | Cephalosporin | **S ($\leq 0.25$)** | **S ($\leq 0.25$)** |
| A | Tilmicosin | Macrolide | **R (> 64)** | **S ($\leq 4$)** |
| A | Tulathromycin | Macrolide | **R (> 64)** | **S (= 4)** |
| A | Penicillin | Penicillin | **R (> 8)** | **S (= 0.25)** |
| A | Florfenicol | Phenicol | **R (> 8)** | **S (= 0.5)** |
| A | Danofloxacin | Fluoroquinolone | **NI (> 1)** | **S ($\leq 0.12$)** |
| A | Enrofloxacin | Fluoroquinolone | **R (> 2)** | **S ($\leq 0.12$)** |
| A | Oxytetracycline | Tetracycline | **R (> 8)** | **S ($\leq 0.5$)** |
| B | Sulfadimethoxine | Sulfonamide | **NI (> 256)** | **NI (> 256)** |
| B | Ampicillin | Penicillin | **NI (> 16)** | **NI ($\leq 0.25$)** |
| C | Tylosin | Macrolide | **NI (> 32)** | **NI (= 32)** |
| D | Gentamicin | Aminoglycoside | **NI (> 16)** | **NI (= 2)** |
| D | Sulfamethoxazole Trimethoprim | Potentiated Sulfa | **NI ($\leq 2$)** | **NI ($\leq 2$)** |
| Ungrouped | Clindamycin | Lincosamide | **NI (> 16)** | **NI (= 8)** |

**R** = Resistant, **S** = Susceptible, **NI** = No interpretation in CLSI group

**Table 2.** 1D base-called read statistics from Albacore for both *Mannheimia haemolytica* strains.

|  | Total Reads | Mean (bp) | Median (bp) | Min (bp) | Max (bp) | N25 | N50 | N75 |
| --- | --- | --- | --- | --- | --- | --- | --- | --- |
| *M. haemolytica* OADDL-1 |  |  |  |  |  |  |  |  |
| FAB 42646 | 27,304 | 7,562 | 5,267 | 210 | 77,585 | 19,083 | 11,974 | 6,444 |
| FAB 42472 | 26,174 | 6,969 | 4,818 | 69 | 65,212 | 17,903 | 10,837 | 5,831 |
| *M. haemolytica* OADDL-2 |  |  |  |  |  |  |  |  |
| FAB 42646 | 14,132 | 11,611 | 9,132 | 241 | 75,460 | 23,851 | 17,018 | 9,985 |
| FAB 42472 | 14,030 | 10,629 | 8,665 | 249 | 68,470 | 22,657 | 15,722 | 8,982 |

### MinION Sequencing

Two MinION R9.4 SpotON flow cells, labeled FAB 42646 and FAB 42472, were used for sequencing. The MinKNOW GUI application (version 1.4.2) was used to perform a platform quality check to ensure flow cell quality. Flow cells were primed as instructed prior to sequencing. MinKNOW version 1.4.2 was executed to sequence using the protocol script NC_48Hr_Sequencing_Run_FLO-MIN105_SQK-LSK208.py. Both libraries were sequenced for approximately 48 hours, with no live base-calling performed.

### Albacore Base-calling

Due to discontinued support of 2-D reads by ONT and Nanopolish (0.6-dev), sequenced reads were subsequently base-called using the newest version of Albacore v 2.1.0. for 1-D base-calling, where both template and complement strands were considered as individual reads.

Albacore performs base-calling on the raw reads using event detection to categorize the signal-level data, after which the event detection is compared to existing models and associated with a nitrogenous base. All raw reads were evaluated and filtered for a cut-off quality score of seven. Base-called reads were further split into their respective barcode folders of *M. haemolytica* OADDL-1 and OADDL-2 for downstream analyses. All reads were trimmed using Porechop (v0.2.1, github.com/rrwick/Porechop) to remove barcoded adapter sequences found in the read files.

### Genome Assembly using Canu

Genome assembly was performed using the software package Canu (version 1.6). Canu implements the overlap-layout-consensus method and is partitioned into three primary stages: correction, trimming and assembly (25). Default Canu command and parameters for assembly were used with a suggested genome size of 2.7 million bp. QUAST was used to evaluate assembly statistics (26).

### Genome Polishing with Nanopolish

Typically, Nanopore sequencing reads are prone to deletion errors, particularly in homopolymer regions (27). In this study, Nanopolish (0.6-dev), a software program that utilizes a trained hidden Markov model to improve the final consensus assembly was used for error correction. Reads were aligned back to the Canu assembly using bwa (28), with the –x ont parameters for noisy ONT reads; samtools was used to sort and index the reads (29). Nanopolish was used to compute a polished consensus genome by correcting substitution, insertion and deletion errors that may have occurred during sequencing or base-calling. Following the github protocol for Nanopolish (https://github.com/jts/nanopolish), the default commands and parameters for computing a consensus sequence were used, with thread count set to four and parallel count set to eight.

### RAST, CARD and ResFinder Annotation

Predicted genes were called from the assembled contigs using the prokaryote gene calling software Prodigal (30). All amino acid sequences were annotated using a combination of homology search with NBCI blast+ (31), and domain identification using hmmscan (32) with PFAM v28.0 database (33). AMR gene determination was performed using NCBI blast+ with three different tools and databases: RAST (Rapid Annotation using Subsystem Technology, (34)), CARD (the Comprehensive Antibiotic Resistance Database, (35, 36)) and ResFinder (37). RAST is a web-based service that offers rapid annotation of prokaryotic genomes (http://rast.nmpdr.org/rast.cgi), using an automated system that requires a genome submission. Similarly, ResFinder is also a web-based service (https://cge.cbs.dtu.dk/services/ResFinder), where AMR genes are identified with the threshold and minimum length set to 90% and 60%, respectively. The CARD database (http://arpcard.mcmaster.ca), which integrates disparate molecular and sequence data, also provides Antibiotic Resistance Ontology (ARO) (36).

### Genome Sub-sampling for AMR Gene Identification

To evaluate the degree to which multiplexity would affect assembly quality for AMR detection, sub-sampling of reads was conducted. Six sub-divisions of randomly sampled reads were created from the total sequencing reads; each subdivision represented 1/10^th^, 1/6^th^, 2/6^th^, 3/6^th^, 4/6^th^ and 5/6^th^ of the total reads. Five replicates were generated for each sub-division of reads; the replications were created to examine the variability derived from random sampling of raw sequencing reads. AMR predictability was then assessed by the quality of the genome assemblies and annotation methods described earlier. For all sub-divided group of reads and replicates, assemblies were generated using Canu v1.6; assembly qualities were evaluated using QUAST and polished with Nanopolish. And, *in silico* determination for AMR genes was conducted by CARD and ResFinder.

## Results

### AMR Phenotyping by MIC

Details of the MIC results are listed in Table 1. Overall, *M. haemolytica* OADDL-1 was found to be susceptible and *M. haemolytica* OADDL-2 was resistant to most of the antimicrobials used in routine therapeutics, with the only exception of OADDL-2 susceptible to Ceftiofur. In most cases for antimicrobials with no CLSI interpretation guidelines (NI), the MIC values for OADDL-2 were found higher than the susceptible OADDL-1; however, both OADDL-1 and OADDL-2 showed similar MIC levels for Sulfadimethoxine and Sulfamethoxazole Trimethoprim (Table 1).

### Sequencing and Base-calling

Initial MinION sequencing yielded 138,689 and 116,073 raw reads for flow cell FAB 42646 and FAB 42472, respectively, for a total of 254,762 reads to be assembled. As shown in Table 2, Albacore 1-D base-calling resulted in 53,478 reads that passed the quality filter for OADDL-1, and 28,162 total reads for OADDL-2. The mean read length for OADDL-1 was 7,562 bp for FAB 42646 and 6,969 bp for FAB 42472 with an N50 value at 6,807 for flow cell FAB 42646 and 8,796 for flow cell FAB 42472. The mean read length for *M. haemolytica* OADDL-2 was 11,611 for FAB 42646 and 10,629 for FAB 42472 and an N50 of 17,018 bp for FAB 42646 and 15,722 bp for FAB 42472 (Table 2).

### Genome Assembly and Genome Polishing

Genome assembly with Canu v1.6 of quality sequencing reads for OADDL-1 resulted in a single contig of 2,644,744 bp with a mean GC content of 41.19%. The draft genome assembly of OADDL-2 was composed of five contigs totaling 2,804,188 bp with an average GC content of 41.24% (Table 3). Polishing (Nanopolish) resulted in increased genome length for both OADDL-1 (2,656,520 bp) and OADDL-2 (2,815,485 bp) (Table 3).

**Table 3.** Assembly statistics generated by Canu and improved by Nanopolish.

|  | Number of contigs | Total length | Total length polished | N50 | GC content (%) |
| --- | --- | --- | --- | --- | --- |
| *M. haemolytica* OADDL-1 | 1 | 2,644,744 bp | 2,656,520 bp | - | 41.19 |
| *M. haemolytica* OADDL-2 | 5 | 2,804,188 bp | 2,815,485 bp | 2,381,899 bp | 41.24 |

The assembly of OADDL-2 also revealed the presence of a 5,265 bp plasmid; a BLAST search returned a 99% identity and 80% coverage of a known plasmid *Pasturella multocida* pCCK411, strain U-B411 (FIG 1); the GC content, and over-and under-representation of GC regions of the plasmid is depicted in FIG 1. A native plasmid found in *M. haemolytica*, plasmid *P. haemolytica,* which encodes for the Bla gene that confers beta-lactam resistance (38) was also found with a BLAST search and returned with a 99% identity and 80% coverage.

**FIG 1.**
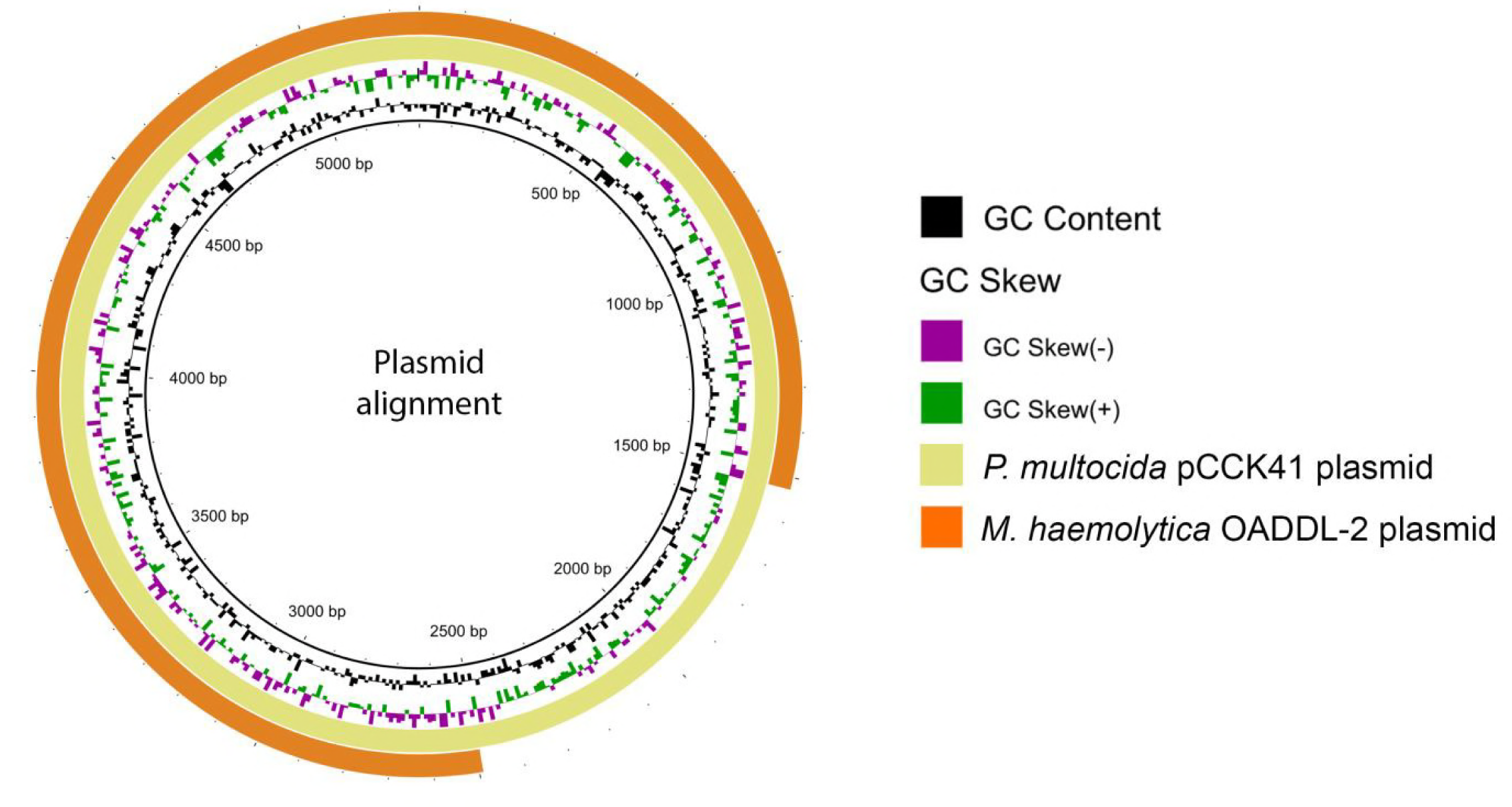
BLAST genome alignment between the plasmid found in *M. haemolytica* OADDL-2 and the highest scored plasmid found in a BLAST alignment plasmid pCCK411. GC-content and GC-skew are also included. GC Skew is shown to visualize regions where there is an over-(+) or under-(-) abundance of guanine and cytosine compared to adenine and thymine.

### Genome Annotation and AMR Gene Identification

Annotation by RAST for OADDL-1 revealed 3,593 coding sequences, which are further categorized under subsystems such as carbohydrates, DNA metabolism, virulence, disease and defense, to name a few. Detailed information of RAST annotations is provided in Table 1S. The RAST annotation for OADDL-2 detected 4,030 coding sequences, of which 45 were related to virulence, disease and defense (Table 2S). Amongst these 4,030 coding sequences, the RAST annotation for OADDL-2 revealed multiple genes that may confer resistance; these include beta-lactamase (bl), Aminoglycoside N6’-acetyltransferase (N6’-ac), and Streptomycin 3”-O-adenylyltransferase (AadA1), for instance (Table 4). The locations of these AMR related genes are showed in FIG. 2.

**Table 4.** Unique AMR genes found in *M. haemolytica* OADDL-2 using the RAST annotation web server.

| Gene Abbreviations | Functional Role |
| --- | --- |
| BL | Beta-lactamase (EC 3.5.2.6) |
| BLc | Beta-lactamase class C and other penicillin binding proteins |
| BLI | Metal-dependent hydrolases of the beta-lactamase superfamily I |
| BLR | Negative regulator of beta-lactamase expression |
| bl | Beta-lactamase |
| BLII | Metal-dependent hydrolases of the beta-lactamase superfamily II |
| BLIII | Metal-dependent hydrolases of the beta-lactamase superfamily III |
| BLA | Beta-lactamase class A |
| CTX | Beta-lactamase CTX-M-16 |
| blaR | Regulatory protein BlaR1 |
| blaI | Beta-lactamase repressor BlaI |
| BLD | Beta-lactamase class D |
| OXA | Beta-lactamase OXA-18 precursor (EC 3.5.2.6) |
| Top | DNA Topoisomerase III (EC 5.99.1.2) |
| AmpS | Beta-lactamase AmpS |
| gyrase | DNA gyrase subunit A and B (EC 5.99.1.3) |
| PSE2 | Beta-lactamase PSE-2 precursor (EC 3.5.2.6) |
| ybxI | Probable beta-lactamase ybxI precursor (EC 3.5.2.6) |
| CS | Beta-lactamase (Cephalosporinase) (EC 3.5.2.6) |
| AadA1 | Streptomycin 3''-O-adenylyltransferase (EC 2.7.7.47) |
| AadA2 | Spectinomycin 9-O-adenylyltransferase |
| N6'-ac | Aminoglycoside N6'-acetyltransferase (EC 2.3.1.82) |

**FIG 2.**
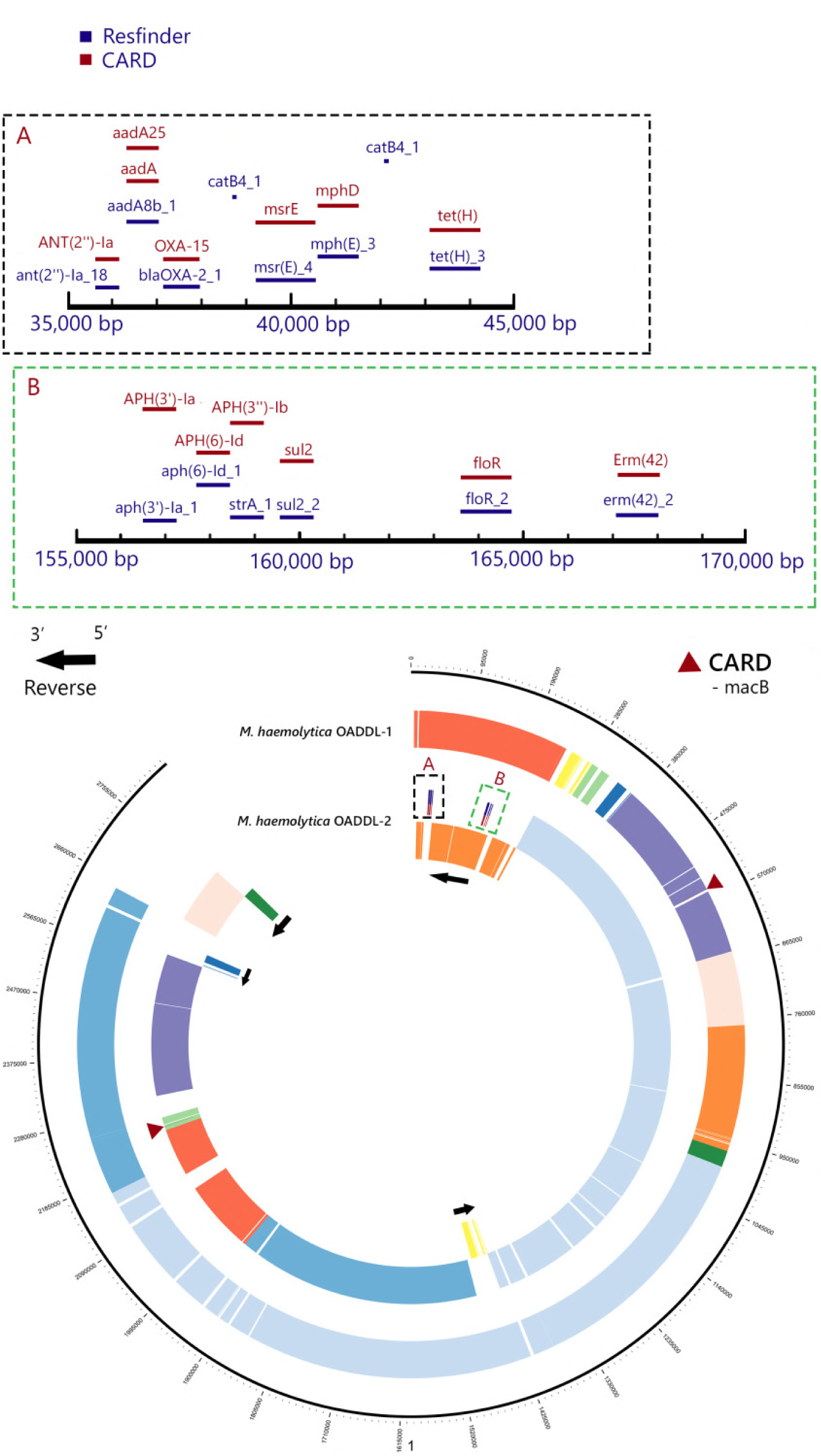
Global genome alignment between bacterial species *M. haemolytica* OADDL-1 and *M. haemolytica* OADDL-2. Homologous genome regions are marked by colors using *M. haemolytica* OADDL-1 as basal frame; the 5’ to 3’ reverse regions in *M. haemolytica* OADDL-2are indicated by the arrows. The *macB* genes, annotated only by CARD, are found in non-homologous regions of *M. haemolytica* (marked by the red triangles). Two genome clusters, one from 35,000 to 45,000 bp (A) and another from 156,000 to 168,000 bp (B) in a reverse region of *M. haemolytica* OADDL-2, accommodate most of the genetics of AMR.

ResFinder, which by default searches resistant genes that share 98% identity to the genes within the databases, was used to identify 18 AMR genes in OADDL-*2* strain (Table 5); examples of these AMR genes are blaOXA-2_1 and blaROB-1_1 Beta-lactam. AMR genes commonly found in *M. haemolytic* resistant strains (21, 39), such as tet(H)_3 (Tetracycline) and aph(3’)-Ia_1 (Aminoglycoside), were also identified in OADDL-2, whereas the susceptible OADDL-1 returned no results for AMR genes on ResFinder.

**Table 5.** Identification of AMR genes with ResFinder and CARD databases.

| AMR Genes | Database | Antibiotic Class | *M. haemolytica*<br>OADDL-2 | *M. haemolytica*<br>OADDL-1 | *M. haemolytica*<br>89010807N | *M. haemolytica*<br>M42548 |
| --- | --- | --- | --- | --- | --- | --- |
| ant(2'')-Ia_18 | ResFinder | Aminoglycoside | Present | - | - | - |
| aadA8b_1 | ResFinder | Aminoglycoside | Present | - | - | - |
| blaOXA-2_1 | ResFinder | Beta-lactam | Present | - | - | - |
| msr(E)_4 | ResFinder | MLSB | Present | - | - | - |
| mph(E)_3 | ResFinder | Macrolide | Present | - | - | - |
| catB4_1 | ResFinder | Phenicol | Present | - | - | - |
| tet(H)_3 | ResFinder | Tetracycline | Present | - | Present | Present |
| blaROB-1_1 | ResFinder | Beta-lactam | Present | - | - | - |
| blaROB-1_1 | ResFinder | Beta-lactam | Present | - | - | - |
| aph(3')-Ia_1 | ResFinder | Aminoglycoside | Present | - | - | Present |
| aph(6)-Id_1 | ResFinder | Aminoglycoside | Present | - | - | Present |
| aph(3'')-Ib_3 | ResFinder | Aminoglycoside | Present | - | - | - |
| sul2_11 | ResFinder | Sulfonamide | Present | - | - | - |
| floR_2 | ResFinder | Phenicol | Present | - | - | - |
| erm(42)_2 | ResFinder | MLSB | Present | - | - | - |
| strA_1 | ResFinder | Aminoglycoside | - | - | - | Present |
| sul2_2 | ResFinder | Aminoglycoside | - | - | - | Present |
| ANT(2'')-Ia | CARD | Aminoglycoside | Present | - | - | - |
| aadA | CARD | Aminoglycoside | Present | - | - | - |
| aadA25 | CARD | Streptomycin/Spectinomycin | Present | - | - | - |
| OXA-15 | CARD | Beta-lactam | Present | - | - | - |
| msrE | CARD | Erythromycin/Streptogramin | Present | - | - | - |
| mphD | CARD | Macrolide | Present | - | - | - |
| tet(H) | CARD | Tetracycline | Present | - | Present | Present |
| ROB-1 | CARD | Beta-lactam | Present | - | - | - |
| macB | CARD | Macrolide | Present | Present | - | - |
| APH(3')-Ia | CARD | Aminoglycoside | Present | - | - | Present |
| APH(6)-Id | CARD | Aminoglycoside | Present | - | - | Present |
| APH(3'')-Ib | CARD | Aminoglycoside | Present | - | - | Present |
| sul2 | CARD | Sulfonamide | Present | - | - | Present |
| floR | CARD | Chloramphenicol | Present | - | - | - |
| Erm(42) | CARD | MLSB | Present | - | - | - |

Alternatively, the AMR gene homology search using CARD identified 16 AMR genes in the OADDL-2 resistant strain; common AMR genes like ROB-1 Beta-lactam, tet(H) for Tetracycline and APH for Aminoglycoside were present in the resistant *M. haemolytica* strain, and two other published strains 89010807N (GenBank accession no. CP011098) (40) and M42548 (GenBank accession no. CP005383) (41) (Table 5). Interestingly, *macB* that confers resistance to macrolide antibiotics was identified for both OADDL-1 and OADDL-2 (Table 5), however not in homologous regions (dark red triangles in FIG 2). This *macB* annotation, though present in both by CARD, showed sequence coverage as low as 10.65% (OADDL-1).

A global alignment to identify homologous genomic regions between OADDL-1 and OADDL-2 was performed using Mauve (42); the aligned regions by Mauve were visualized using Circos (43), with the annotations for AMR genes by both CARD and ResFinder. In FIG 2, regions with shared colors indicate sequence homology. Though the overall sequence similarity was estimated at 95.2%; significant structural variation is also identified, including four reverse regions in OADDL-2 (FIG 2). Among these reverse regions, two genome clusters in the orange colored region of OADDL-2, homologous to the 760,000 to 950,000 bp region of the susceptible OADDL-1, accommodate most of the annotated AMR related genes (details in the A and B boxes of FIG 2).

### Genome Assembly Quality Assessment and AMR Gene Identification

To assess adequate amounts of long reads for microbial genome assembly and subsequent AMR gene detection, subsampling was conducted at 1/10^th^, and from 1/6^th^ to 5/6^th^ of the total reads used for original assembly. In OADDL-1, the base pair length was consistent across all replicates and partitions (mean=2,649,246 bp), with standard deviations ranging from 9,575 bp (1/6^th^) to 36,605 bp (1/10^th^). Mean GC content was also stable across all samples with a range of 41.16% to 41.23% (Table 6). Assembly quality began to decline at 2/6^th^ of the original total reads (8,913 reads); average N50 reduced to 1,864,819 bp, and averaged number of mismatches with original assembly increased to 17,233 bp. At 1/10^th^ of the total reads, the number of contigs increased 10 fold (19 to 24 contigs) and the N50 of the assemblies dropped to 351,563 bp, showing a significantly reduced assembly quality compared to the other subsamples.

**Table 6.** Assembly quality assessment of sequencing read subsets; results of *M. haemolytica* OADDL-1.

| Subsample | Average base-pair length<br>(std. dev.) | Contig count | Average N50<br>(std. dev.) | GC Content<br>(%) | Ave. Tot. Mismatches<br>(std. dev.) |
| --- | --- | --- | --- | --- | --- |
| 1/10<sup>th</sup> of reads | 2,628,537 (36,605) | 19 - 24 | 351,563 (224,531) | 41.2 – 41.3 | 30,993.6 (531.6) |
| 1/6<sup>th</sup> of reads | 2,624,083 (9,575) | 4 - 10 | 1,292,922 (518,613) | 41.2 – 41.2 | 20,988.0 (131.9) |
| 2/6<sup>th</sup> of reads | 2,668,633 (13,312) | 2 - 5 | 1,864,819 (652,239) | 41.2 – 41.2 | 17,233.4 (231.7) |
| 3/6<sup>th</sup> of reads | 2,662,841 (13,994) | 1 - 4 | 2,610,430 (41,359) | 41.2 – 41.2 | 15,802.6 (365.1) |
| 4/6<sup>th</sup> of reads | 2,660,579 (14,754) | 1 - 4 | 2,576,912 (71,524) | 41.2 – 41.2 | 14,632.6 (185.2) |
| 5/6<sup>th</sup> of reads | 2,650,805 (16,078) | 1 - 3 | 2,589,962 (111,922) | 41.2 – 41.2 | 13,939.0 (181.4) |
| Mean | 2,649,246 (25,931) | - | - | - | 18,931.5 (271.2) |

For OADDL-2, subsampling of the reads generated comparable results. The average genome size of all subsampled assemblies was 2,777,982 bp, with a standard deviation of 24,724 bp; mean GC content remained consistent amongst these assemblies with a range of 41.22% to 41.33% (Table 7). The plasmid was found in all assemblies. Partitioning of sequencing reads at 1/10^th^ showed a rapid degradation in assembly quality: the number of contigs increased to at least 29 (maximum at 40), and the N50 value dropped to an average of 133,657 bp (Table 7).

**Table 7.** Assembly quality assessment of sequencing read subsets and AMR gene identification; results of *M. haemolytica* OADDL-2.

| Subsamples | Average base-pair<br>length (std. dev.) | Contig<br>count | Average N50<br>(std. dev.) | GC Content<br>(%) | Plasmid<br>(Y/N) | AMR genes<br>ResFinder (mean) | AMR genes<br>CARD (mean) | Ave. Tot. Mismatches<br>(std. dev.) |
| --- | --- | --- | --- | --- | --- | --- | --- | --- |
| 1/10<sup>th</sup> of reads | 2,619,204 (43,898) | 29 - 40 | 133,657 (9,800) | 41.2 - 41.3 | Y | 15-17 (15.6) | 14-16 (15) | 37,875.2 (599.8) |
| 1/6<sup>th</sup> of reads | 2,800,552 (16,129) | 6 - 20 | 606,852 (471,233) | 41.3 - 41.3 | Y | 15-17 (15.6) | 14-16 (15) | 25,032.6 (559.5) |
| 2/6<sup>th</sup> of reads | 2,806,768 (16,792) | 4 - 6 | 2,178,862 (407,453) | 41.2 - 41.3 | Y | 15-16 (15.4) | 14-15 (14.6) | 17,772.2 (292.8) |
| 3/6<sup>th</sup> of reads | 2,792,790 (14,509) | 3 - 6 | 2,197,805 (532,021) | 41.2 - 41.3 | Y | 15-17 (15.6) | 14-15 (14.6) | 15,962.6 (145.8) |
| 4/6<sup>th</sup> of reads | 2,844,871 (40,342) | 3 - 8 | 2,254,699 (412,158) | 41.2 - 41.3 | Y | 14-16 (15) | 14-15 (14.6) | 14,807.2 (221.7) |
| 5/6<sup>th</sup> of reads | 2,803,710 (16,678) | 3 - 4 | 2,602,409 (349,320) | 41.3 - 41.3 | Y | 15-16 (15.2) | 14-15 (14.4) | 14,064.8 (173.2) |
| Mean | 2,777,982 (24,724) | - | - | - | - | 15.4 | 14.7 | 20,919.1 (332.1) |

For all assemblies using subsets of reads, annotation was done by ResFinder and CARD databases. Overall, there were no observable changes compared to the original annotation for AMR genes found in OADDL-2. CARD AMR annotation varied slightly due to the low sequence identity of the *macB* gene. With the exception of the *macB* gene present at low coverage, there were no resistance genes found on OADDL-1.

## Discussion

Massive parallelization of sequencing reaction, such as the popular Illumina platforms (44), has remarkably increased the read numbers per run to a few billion of short read fragments, transforming DNA molecular readers into diagnostic tools in virtually every field of biomedical research (45-47). Among all available technologies, growing interest was recognized in the field of long-read sequencing, owing to the apparent genome complexity in lengthy repetitive elements, copy number and structural variations that lead to difficulty in assembling data from short-read platforms (44). Currently, the dominant platforms for direct typing long read fragments are Pacific Biosciences (PacBio) and Oxford Nanopore, both employing a single-molecule real-time approach. In addition to the improved assembly when kilobase sized reads are available (48), single molecular technologies are amplification free and thereby not hampered by PCR-based artifacts and GC-bias, giving a much more uniform coverage and span across GC-rich regions (49). The feasibility of ONT has been studied in clinical plasmid isolates of *Escherichia coli* (14), *Salmonella typhimurium* (15), *Vibrio parahaemolyticus* (15) and *Klebsiella pneumoniae* (14, 15). Nanopore-only assembly yielded > 99% (0.2% mismatches and 0.5-0.6% gaps) consensus accuracy compared to Illumina MiSeq, and with polishing using MiSeq paired-end reads, assembly accuracy increased to ∼ 99.9% (14). However, when multiplexing 12 plasmids by Oxford’s Rapid Barcoding Sequencing kit, the average accuracy of two plasmids compared to the known reference assembly turned out to be considerably lower (87% in (15)). In Wick et al. (2017), 12 isolates of *K. pneumonia* were barcoded, multiplexed and sequenced with 1-D technology on a single flow cell; the most accurate ONT-only assembly had an estimated 0.349% error rate after polishing (50). These results have recommended an alternative approach of using both Illumina and ONT reads for hybrid assemblies (51).

To study AMR detection capacity of ONT, we sampled reads from MinION R9.4 flow-cells. When using the full capacity, one *M. haemolytica* strain was fully assembled while the other produced 5 contigs with N50 that covers over 85% of the polished genome. The averaged genome coverage was at ∼146x and ∼111x for OADDL-1 and OADDL-2, respectively; both assemblies showed a GC content (Table 3) similar to other published strains (∼ 41% in (40, 41, 52, 53)). Results from our barcoded, multiplexed *M. haemolytica* strains suggest that consistent taxonomic status and genome evolvability and transduction capacity could be correctly determined (54-56). Capacity of ONT’s single molecule long-read sequencing platform for strain identification is thus confirmed.

In this study, AMR profiles of both *M. haemolytica* strains were determined by standard MIC method (results see Table 1). In CLSI guidelines (24), drugs with host specific MIC interpretation are categorized as Group A drugs. Several of the Group A drugs listed in Table 1 are commonly used to treat bacterial respiratory disease in cattle. However for cases like Danofloxacin, though it is classified as Group A, bovine specific MIC breakpoints for resistance has yet not been defined, making accurate interpretations of resistant phenotypes ambiguous. For drugs not classified under Group A, interpretations of MIC values impose greater challenges, since they must be predicted based on guidelines for human or other host species. From a public health perspective it is significant to understand the resistance mechanisms for all groups of antimicrobials whether they are used in animals or not. Direct identification of the genomic basis for AMR, such as our study case in *M. haemolytica*, could be a more reliable approach.

We also examined different aspects that could impact the cost-effectiveness for broader applicability of ONT to for AMR detection. Aside from the issues associated with the biased input of barcoded DNA samples (50), at a greater degree of multiplexity assembly quality could be concerning, which might subsequently lead to mis-identification of AMR genes. Current ONT R9.4.1 flow cell technology with SQK-LSK 109 sequencing kits averages over 10 billion bp per flow cell. When genome coverage and assembly quality are not the major concerns, our results indicate that rapid AMR diagnosis using ONT long-read sequencing technology could provide sufficient capacity for genotyping over 12 microbial genomes simultaneously in one ONT flow cell in 48 hours. The limitation, in principle, is due to the use of ONT barcoding kit (EXP-NBD103), which works in conjunction with the current genomic DNA sequencing kit SQK-LSK109. Here we also should note that, unlike Wick et al. (2017) (50), our simulated genome assemblies listed in Table 6 and Table 7 were assembled using ONT-only reads, but polished on the constructed genome of OADDL-1 with all reads (17,714 reads; 244 million bp). There are other proven cases that demonstrate how superior assembly quality can be obtained through hybrid assembly by ONT-Ilumina (57, 58) and PacBio-Illumina (59, 60). Based on our ONT-only read assemblies, AMR genes identified in *M. haemolytica* strains were congruous to the MIC results (Table 1). Amongst the tested antibiotics, Ceftiofur, a third generation of cephalosporin antibiotics as well as a beta-lactam, was found to be susceptible in both strains of *M. haemolytica* (Table 1). It has been suggested that resistance to broad spectrum cephalosporins such as Ceftiofur is conferred by AmpC type beta-lactamases (61, 62), which were not detected in either of the *M. haemolytica* isolates used in this study. Studies focusing on comparisons of phenotypic susceptibility testing with sequence-based AMR prediction have already been performed for a variety of bacterial pathogens (8, 63-65), now to include this report on *Mannheimia* strains. Amongst these, promising results of 97% sensitivity and 99% specificity for AMR detection was reported from a large study conducted in U.K. for *Staphylococcus aureus* strains (8), clearly demonstrating that WGS is transforming the practice of outbreak investigation with remarkable accuracy.

Equipped with such sequencing capacity, we believe direct genome sequencing approaches for AMR detection has started to shed light on genomic surveillance for disease control. At present, routine drug susceptibility testing is undertaken using phenotype-based methods, including disc diffusion, gradient diffusion and broth dilution methods that have been automated in a number of commercial platforms (64, 66). Despite the problems with specificity for certain organism-antimicrobial combinations (67, 68) and common operation errors in practices and result interpretation (69), drug susceptibility testing requires laboratory procedures that could be completed within 48 ∼ 96 hours. Further, standard susceptibility testing protocols can only be done for culturable bacteria, creating significant challenges for risk assessment when the entire AMR reservoir needs to be investigated for both culturable and unculturable pathogens (70). Bacterial unculturability is therefore the most problematic for complete AMR detection, as it is possible that only about 1% of what is present in a sample can be grown with current cultivation techniques (71, 72). Determination of AMR of anaerobic and other difficult to grow fastidious bacteria is also challenging. Genomic approaches (73, 74), on the other hand, can provide sufficient information to re-assemble genomics of AMR for ‘unculturable’ samples (12, 73). To fulfill the need for timely response in disease control, WGS approaches that rely on growing, isolating and purifying bacterial isolates might still fall short. To effectively reduce the response time and expand genomic surveillance for “unculturable” pathogens, direct sequencing approaches should be considered. This culture-independent approach is encouraged by the recent successes in meta-genomics (sequencing all genomes in a sample) (74-76), though much still remains to be studied, like contaminants and low biomass specimens, prior to the broad adoption of meta-genomics in diagnostic laboratories (77).

## Accession number(s)

The MinION data obtained in this study have been deposited in the NCBI Sequence Read Archive under BioProject PRJNAxxxxxx. (please contact for early access).

## Acknowledgements

Funding for this study was provided for A.R., by the Center for Veterinary Health Sciences, OSU. This research is also supported by the NSF-MRI 1626257 for C.C.; the work presented in this report reflects the support from the USDA HATCH project OKL03011 of C.C.. Data analysis was completed with support from the High Performance Computing Center Facilities at Oklahoma State University.

